# Selective K-ATP channel-dependent loss of pacemaking in vulnerable nigrostriatal dopamine neurons by α-synuclein aggregates

**DOI:** 10.1101/842344

**Authors:** Poonam Thakur, Kelvin Luk, Jochen Roeper

**Author notes:** corresponding author Dr. Poonam Thakur, Institute for Neurophysiology, Neuroscience Center, Goethe University, Frankfurt, Germany.

## Abstract

Parkinson disease (PD), one of the most common neurodegenerative disorder, is believed to be driven by toxic α-synuclein aggregates eventually resulting in selective loss of vulnerable neuron populations, prominent among them, nigrostriatal dopamine (DA) neurons in the lateral substantia nigra (l-SN). How α-synuclein aggregates initiate a pathophysiological cascade selectively in vulnerable neurons is still unclear. Here, we show that the exposure to low nanomolar concentrations of α-synuclein aggregates (i.e. fibrils) but not its monomeric forms acutely and selectively disrupted the electrical pacemaker function of the DA subpopulation most vulnerable in PD. This implies that only dorsolateral striatum projecting l-SN DA neurons were electrically silenced by α-synuclein aggregates, while the activity of neither neighboring DA neurons in medial SN projecting to dorsomedial striatum nor mesolimbic DA neurons in the ventral tegmental area (VTA) were affected. Moreover, we demonstrate functional K-ATP channels comprised of Kir6.2 subunit in DA neurons to be necessary to mediate this acute pacemaker disruption by α-synuclein aggregates. Our study thus identifies a molecularly defined target that quickly translates the presence of α-synuclein aggregates into an immediate impairment of essential neuronal function. This constitutes a novel candidate process how a protein-aggregation-driven sequence in PD is initiated that might eventually lead to selective neurodegeneration.

## INTRODUCTION

α-Synuclein is a small 14 kDa protein located mainly in presynaptic terminals, soma and nucleus (Maroteaux et al., 1988). While its physiological function is still unclear, its causal role in the pathogenesis of Parkinson disease (PD) is well established (Polymeropoulos et al., 1997; Simón-Sánchez et al., 2009). Occurrence of toxic gain-of-function α-synuclein aggregates and their spreading within various regions of central and peripheral nervous system has been suggested to be intimately linked to disease progression (Spillantini and Crowther, 1998). Aggregated α-synuclein has amyloid conformation rich in beta sheet structure (Roeters et al., 2017), different from the natively unfolded conformation of monomeric forms (Eliezer et al., 2001). Furthermore, various synucleinopathies are also believed to act via unique conformers of α-synuclein (Peng et al., 2018). Various cell-specific toxicity profiles of α-synuclein are determined closely by its exact conformation or strain (Bousset et al., 2013; Peelaerts et al., 2015). These fibrillar forms of α-synuclein can be produced in vitro and, upon application, are quickly internalized by cells, where they seed the aggregation of native monomeric α-synuclein in a prion like fashion. (Volpicelli-Daley et al., 2014).

While α-synuclein rich Lewy bodies are observed in numerous brain regions (Braak et al., 2004), all affected neurons are not equally susceptible to degeneration. It is well known that DA neurons of substantia nigra (SN) are a major – and clinically most relevant - target neuronal population for neurodegeneration during PD and leads to the motor symptoms of disease (Greffard et al., 2006). DA neurons in the neighboring ventral tegmental area (VTA) region, however do not show degeneration and are much more resistant to toxic insults in animal models (Thakur et al., 2017). Even within SN there are regional variations in patterns of cell loss, e.g. ventrolateral tier shows a much bigger and faster loss of neurons during disease course in comparison to dorsomedial tier (Kordower et al., 2013a).

It is, however, not yet clear how α-synuclein aggregates, once present in vulnerable neurons, prominently among them nigrostriatal DA neurons, initiate a cascade of pathophysiological events that eventually lead to the well-established pattern of selective neurodegeneration. “High vulnerability” of DA neurons has previously been characterized by particular functional or structural features – such as high levels of intrinsic redox-stress (Dryanovski et al., 2013), large activity-dependent energy demands or maintenance of an extensive axonal morphology (Matsuda et al., 2009; Bolam and Pissadaki, 2012). A prominent factor might also be a particular style of apparently “dangerous & capricious” calcium handling in the sense of a low capacity for calcium (Ca^2+^) buffering (i.e. most vulnerable DA SN neurons do not express calbindin (Liang et al., 1996)) combined with a high activity-dependent calcium loading of cytosolic and mitochondrial compartments, which is driven by several types of voltage-dependent calcium channels (Chan et al., 2007; Guzman et al., 2009; Ortner et al., 2017; Surmeier et al., 2017; Benkert et al., 2019; Liss and Striessnig, 2019).While the vulnerable SN DA neurons rely on voltage-gated Ca^2+^ channels for generation of subthreshold membrane potential oscillations to drive the pacemaker firing activity (Puopolo et al., 2007), neighboring VTA DAergic neurons utilize tetrodotoxin-sensitive sodium channels (Khaliq and Bean, 2010) and hyperpolarization-activated cyclic nucleotide-gated cation (HCN) channels (Neuhoff et al., 2002). In addition, several types of voltage gated Ca^2+^ channels in SN neurons also contribute to a large influx of Ca^2+^ ions with each action potential (Hage and Khaliq, 2015; Philippart et al., 2016; Ortner et al., 2017).

This activity dependent Ca^2+^ entry into the neurons with each action potential, leads to increase in cellular oxidative stress particularly in mitochondria (Guzman et al., 2010). Ca^2+^ can interact with other cellular stressors such as α-synuclein to cause a selective increase in vulnerability of SN DA neurons but not of the neighboring VTA neurons (Lieberman et al., 2017). Further, only 35% of SN DA neurons express calbindin in contrast to 58% of VTA DA neurons (Dopeso-Reyes et al., 2014). These calbindin positive neurons have been shown to be more resistant to degeneration in several types of animal models of PD (Liang et al., 1996; Kim et al., 2000; Dopeso-Reyes et al., 2014). Increased fraction of calbindin positive neurons among surviving DA neurons might point to a central role of the coupling between high Ca^2+^ influx and metabolic stress in driving PD etiology.

Ca^2+^ entering SN DA neurons might also lead to activation of K-ATP channels thereby facilitating bursting in medial SN DA neurons (Knowlton et al., 2018). We have previously shown that K-ATP channels, which are composed of inward rectifying K channel subunits (Kir6.2) and sulfonylurea receptors subunits (SUR1) in DA neurons mediate SN DA vulnerability in PD models (Liss et al., 2005) and are necessary for in vivo burst firing in medial DA SN neurons (Schiemann et al., 2012). However, their functional and potential pathophysiological role in most vulnerable, lateral SN DA neurons is still unknown.

In addition, we have previously shown that mild overexpression of mutant A53T-α-synuclein lead to in vivo hyperactivity of DA SN neurons via oxidative impairment of Kv4.3 channels (Subramaniam et al., 2014). However, this study focused mostly on the medial SN, where this selective hyperexcitability phenotype of DA SN neurons took more than 6-month in vivo to develop. This study also did not address the question which particular conformer of α-synuclein was responsible. In general, other studies demonstrated that oligomeric/fibrillar α-synuclein is more detrimental to cellular functions in comparison to monomeric α-synuclein, also in DA neurons. For instance, exposure of cultured DA neurons to 100 nM oligomeric α-synuclein for 5-10 minutes was shown to trigger the opening of permeability transition pore and increase oxidative modifications of ATP synthase beta subunit in mitochondria (Ludtmann et al., 2018). This was in contrast to the monomeric form that increased the activity of ATP synthase to improve the efficiency of ATP production (Ludtmann et al., 2016).

In summary, it is still unknown whether α-synuclein aggregates induce acute and/or selective functional changes in the most vulnerable, lateral SN DA neurons. In the present study, we demonstrate that these dorso-lateral striatum projecting lateral SN (DLS-lSN) DA neurons respond with an acute disruption of pacemaker firing when challenged with low nanomolar concentrations of α-synuclein fibrils. In contrast, dorso-medial striatum projecting medial SN (DMS-mSN) DA neurons and nucleus accumbens core projecting DA neurons in VTA (NAcC-VTA) maintained their pacemaker function in response to α-synuclein fibrils. Finally, we demonstrate that selective activation of K-ATP channels in DLS-lSN neurons is necessary to mediate this effect of α-synuclein fibrils on electrical activity.

## MATERIAL AND METHODS

### Identification of various dopamine neuron sub-populations by retrograde tracing

All the experiments were performed on male C57Bl/6N (Charles Rivers) mice aged 8-10 week. Red fluorescent retrobeads (Lumafluor) were diluted (1:30) in ACSF and stereotaxically injected into brain. Mice were anesthetized by exposure to 4% isoflurane in a closed chamber and maintained at 2-2.5 % level during the surgery. 100 nanoliters of diluted beads were injected bilaterally in either dorso-medial striatum (DMS) (AP-0.74, ML-1.1, DV-2.6); dorso-lateral striatum (DLS) (AP-0.74, ML-2.2, DV-2.6) or nucleus accumbens core (NAcC) (AP-1.54, ML-1.0, DV-4). AP coordinates were adjusted to skull size according to the equation: Final AP coordinate = AP coordinate/4.2*distance (bregma - lambda) + 0.3. Period needed for sufficient labelling of target regions was 3 days for DMS and NAcC, and 2 days for DLS. Examples of injection sites in NAcC, DMS and DLS are shown in figure 1(a).

**Figure 1.**
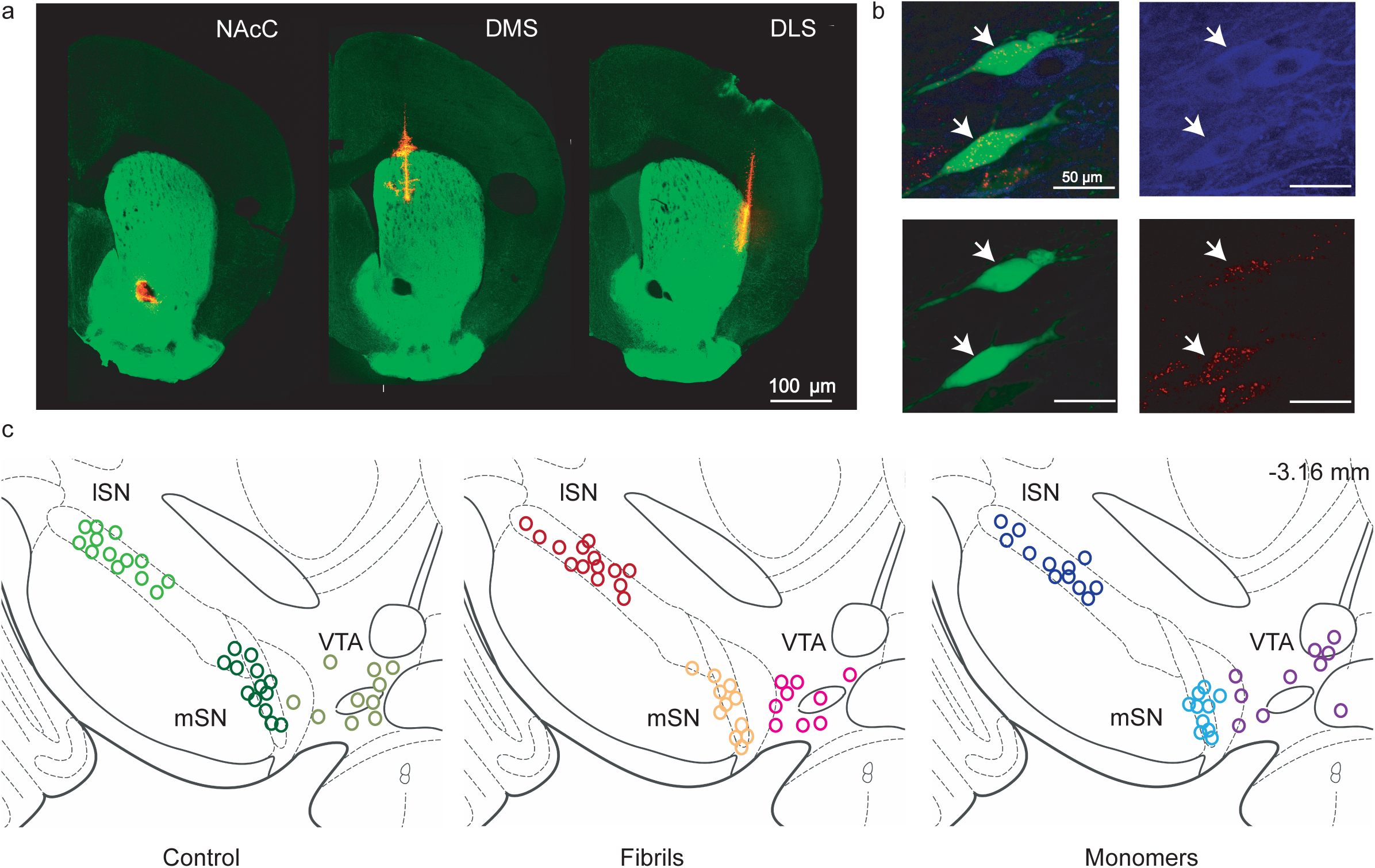
Anatomical mapping of the recorded neurons. **a** Representative images of injection sites in Nucleus Accumbens Core (NAcC), Dorso-Medial Striatum (DMS) and Dorso-Lateral Striatum (DLS) with TH in green and fluorescent retro-beads in red. **b** Representative confocal image of retrogradely labelled neurons (marked with arrows) filled with neurobiotin during the recordings. Green-Neurobiotin, Blue-TH, Red-Retrobeads. **c** Anatomical mapping of all the recorded neurons shown in figure 2, 3 and 4. Each panel shows neurons located in lateral substantia nigra (lSN), medial substantia nigra (mSN) and ventral tegmental area (VTA) in each treatment group-control, fibrils and monomer.

### Electrophysiological patch-clamp recordings

Mice were perfused with ice-cold ACSF (50 mM sucrose, 125 mM NaCl, 2.5 mM KCl, 25 mM NaHCO_3_, 1.25 mM NaH_2_PO_4_, 2.5 mM glucose, 6.2 mM MgCl_2_, 0.1 mM CaCl_2_ and 2.96 mM kynurenic acid (Sigma), oxygenated with 95%O_2_/5%CO_2_). Mid-brain slices were cut coronally using a vibrating microtome at a thickness of 250 µm and allowed to recover for 50 minutes in oxygenated ACSF (22.5 mM sucrose, 125 mM NaCl, 2.5 mM KCl, 25 mM NaHCO_3_, 1.25 mM NaH_2_PO_4_, 2.5 mM glucose, 2.1 mM MgCl_2_ and 2 mM CaCl_2_; 95% O_2_/5% CO_2_) at 37 °C. Recording chamber was continuously perfused with ACSF at a flow rate of 2–4 ml/min. Synaptic input was blocked by addition of CNQX (12.5 µM; Biotrend), D-AP5 (10 µM; Biotrend) and gabazine (SR95531, 4 µM; Biotrend).

SN or VTA DA neurons were visualized by infrared differential interference contrast video microscopy. Retrobeads were visualized by epifluorescence (Zeitz, Jena, Germany). Recording pipettes (4-5 MΩ) were fabricated from thin walled borosilicate glass (GC150TF; Harvard Apparatus, Holliston, MA, USA) and filled with internal solution containing 135 mM K-gluconate, 5 mM KCl, 10 mM HEPES, 0.1 mM EGTA, 2 mM MgCl_2_, 2 mM MgATP, 0.2 mM NaGTP, and 0.1 % neurobiotin, pH 7.4 (285-290 mOsm). Recordings were performed in current clamp or voltage-clamp mode using an EPC-10 patch amplifier (HEKA) with a sampling rate of 10 kHz (low-pass filter, 5 kHz). Electrophysiological data analysis was carried out using custom written scripts in MATLAB_R2017a.

DA neurons containing retrobeads were first recorded in the cell-attached mode to ensure intact pacemaking activity for at least 30 seconds. Afterwards, whole cell configuration was obtained by applying a mild suction. Only neurons with uncompensated series resistance < 10 mOhm were used for analysis. After break in to the whole-cell configuration, recordings in continuous current-clamp were made for a duration of at least 300 seconds to monitor changes in pacemaker function (recordings were carefully terminated via slow conversion to the out-out configuration, which lead to a high success rate (> 95%) of recovering bead-positive, neurobiotin & TH-immunopositive (i.e. DA) neurons post-hoc.

For the extracellular application, 10 nM fibrils were prepared in extracellular solution (0.5 M NaCl, 10 mM HEPES buffer, pH 7.49). Recording pipettes (9-10 MΩ) were fabricated from thick walled borosilicate glass (GC150TF; Harvard Apparatus, Holliston, MA, USA and filled with regular internal solution as described before. After giga-seals were obtained, pacemaker firing was recorded in the voltage clamp mode without breaking into the cell. Once regular firing was observed for at least 30 seconds, extracellular solution (either control or with 10 nM fibrils) was puffed on the cell being recorded using a second pipette with mild pressure (30-50 mbar) so that seal was not disturbed. Recordings were continued for at least 30 seconds post-application of control or 10 nM fibril containing extracellular solution.

After recording, patch slices were post-fixed in 4% paraformaldehyde in PBS, pH 7.4. Striatal regions containing injection sites for retrobeads were also post-fixed and cut at 50 µm thickness for IHC.

### Intracellular dialysis of monomeric and fibrillar α-synuclein into DA neurons

Recombinant human wild type α-synuclein monomeric protein was purified from *E.coli* and assembled into fibrils at 5mg/mL in PBS (pH-7.0) for 7 days as described previously (Volpicelli-Daley et al., 2014). Fibrillization was confirmed using multiple methods (Fig. S1). Aliquots of each reaction tube were centrifuged (45,000 x g, 30 minutes at 23°C) and separated on a 15% SDS gel. Gel was then stained with Coomassie Brilliant Blue and scanned by LiCor (Fig. S1a). The same samples were also assessed using Thioflavin T absorbance assay to confirm amyloid formation (Fig. S1b). To verify their pathogenic potential, 200 nM fibrils were added to mouse hippocampal neurons from embryonic day 16-18 CD1 mice. Neurons were cultured for seven days in poly-D-lysine coated 96-well plates prior to transduction with fibrils and were fixed twelve days post-transduction and stained with 81A antibody to assess the presence of aggregates (Fig. S1c). For electrophysiology experiments, fibrils and monomers were dissolved in sterile dPBS to a stock concentration of 1.25 µg/µl (89.3 µM). 5 µl of this stock solution was diluted with 887 µl of dPBS to obtain an intermediate concentration of 0.5 µM and stored at −80°C. On the day of experiment, to obtain a final concentration of 10 nM, 20 µl of intermediate stock was diluted with 980 µl internal pipette solution (see above) to obtain a final volume of 1 ml. The final solution was sonicated using a Q-sonica probe sonicator (20% power) for 1 minute in a pulsed manner to break down the bigger aggregates of α-synuclein. Monomeric α-synuclein was also prepared in a similar manner but was not sonicated. Control pipette solution was diluted with equivalent amount of dPBS. Final dilution step for both fibrillar and monomeric α-synuclein was carried out immediately before experiment and made fresh for each day of experiment. This approach allowed a targeted application of α-synuclein in each cell.

It should be noted that the molarity calculation is made using the molecular weight of 14 kDa for monomeric α-synuclein. Real molarity of fibrillar solution is expected to be lower than this because of higher molecular weight. However, sonication step leads to a heterogenous mixture with filament sizes usually ranging from 30 nm to 100nm as described previously (Luk et al., 2012; Polinski et al., 2018). Therefore, it is not possible to ascertain the exact molecular weight and molarity of this solution. Addition of monomeric α-synuclein did not cause any change in osmolarity of internal solution (likely due to extremely small concentration). Osmolarity of fibrillar α-synuclein solution could not be measured as the instrument is unable to detect osmolarity in a self-aggregating solution.

### Immunohistochemistry and confocal analysis

Free floating sections were rinsed in PBS and incubated in blocking solution (0.2 M PBS with 10% horse serum, 0.5% Triton X-100, 0.2% BSA). Sections were incubated overnight (room temperature) with goat anti tyrosine hydroxylase (1:1000, Calbiochem/ Merck) antibody diluted in the carrier solution (0.2 M PBS with 1% horse serum, 0.5% Triton X-100, 0.2% BSA). After rinsing with PBS, sections were incubated with goat anti-rabbit Alexa Flour 405 (1:750, Invitrogen). To detect the neurobiotin, streptavidin AlexaFluor-488 (1:1000, Invitrogen) was used. After final rinsing with PBS, sections were mounted on slides with fluorescent mounting medium (Vectashield, Vector Laboratories), coverslipped and visualized with confocal microscope (Nikon Eclipse90i, Nikon GmbH). Location of each recorded neuron was documented and manually reconstructed using the mouse brain atlas (Paxinos and Watson, 2007) as a reference map.

### Statistical analysis

All the statistical tests were performed using GraphPad Prism 7 (GraphPad Software, Inc., La Jolla, CA, USA). Values are given as mean ± sem. One-way ANOVA with Holm Sidak’s post-hoc correction was applied to compare the groups for all analysis unless specified. P values less than 0.05 were considered significant.

## RESULTS

### Nanomolar alpha-synuclein fibrils but not monomers disrupt pacemaker firing of vulnerable dopamine neurons

All the recorded midbrain neurons were identified based on their axon projection targets in dorsal or ventral striatum and were immuno-positive for TH (i.e. were DAergic). Figure 1a shows the representative injection sites for fluorescent retro beads in distinct regions of striatum and nucleus accumbens. Neurons were selected for electrophysiological recordings based on the presence of fluorescent beads and filled with neurobiotin during the recording. As shown in Figure 1b, expression of TH was documented in neurobiotin- and beads-labelled cells. Anatomical location of all recorded neurons was mapped according to the mouse brain atlas (Fig 1c).

Figure 2 shows representative traces of whole-cell patch-clamp recordings of DLS-lSN DA neurons firing under control conditions (Fig. 2a) and after exposure to 10 nM α-synuclein fibrils (Fig. 2b) or 10 nM α-synuclein monomers (Fig. 2c). In the presence of synaptic blockers, we monitored spontaneous activity in the on-cell mode to exclude recording from DA neurons damaged by the slice preparation. Shortly after breaking into the whole cell configuration, we switched to current-clamp and monitored spontaneous electrical activity for at least 300 seconds. Figure 2 shows the effect of dialyzing 10 nM of α-synuclein fibrils into a DLS-lSN DA neuron via the patch pipette. In contrast to recordings of DLS-lSN DA neurons with control pipette solutions, which showed stable (with some degree of slowing), regular and continuous pacemaking, DLS-lSN neurons that were patched with pipette solution containing 10 nM α-synuclein fibrils displayed a dramatic, about 90% decrease in pacemaker frequency within 5 minutes (Fig. 2d). Overall, there was a significant (p< 0.01) decline in steady state firing rate (i.e. firing rate in last 10 seconds of recording) in neurons dialyzed with fibrils in comparison to neurons filled with control recording solution.

**Figure 2.**
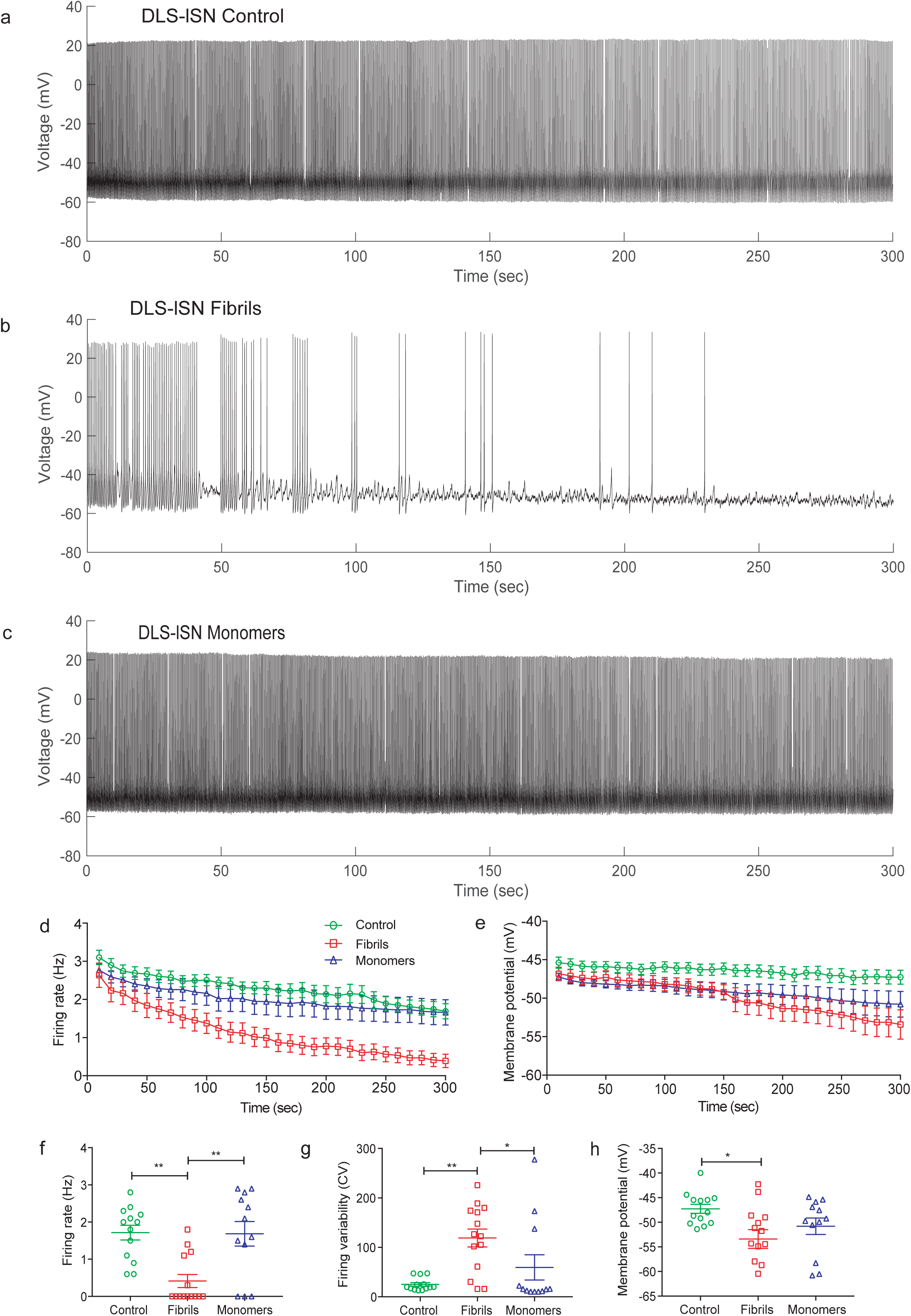
Acute exposure to α-synuclein fibrils disrupts the pacemaker firing of DLS-lSN neurons. Representative trace of whole cell recordings for a duration of 300 seconds from DLS-lSN under **a** control conditions showing regular firing **b** with 10 nM fibrils in the recording solution where firing becomes irregular and stops eventually **c** with 10 nM monomers in the recording solution showing no change in firing activity. Graph shows the change in **d** average firing frequency and **e** average membrane potential over the recording duration in various groups. **f** Comparison of steady state firing frequency (i.e. frequency during last 10 seconds of recording) shows a significant decline (p< 0.001, Ordinary one-way ANOVA followed by Holm Sidak’s multiple comparison test) in firing frequency after fibrils wash-in in comparison to control but not after monomers wash-in. **g** Comparison of regularity of firing (measured as coefficient of variation of inter-spike interval) show a drastic increase in irregularity after fibrils wash-in but not in monomers when compared to control. **h** Comparison of steady state membrane potential (i.e. membrane potential during last 10 seconds of recording) also shows a significant decline (Ordinary one-way ANOVA followed by Holm Sidak’s multiple comparison test) in membrane potential after fibrils wash-in in comparison to control. All data is represented as mean ± sem. Control- n=13, N=9; Fibrils- n=14, N=3; Monomers- n=12, N=3, where n= number of neurons, N= number of mouse.

The frequency reduction was accompanied by a significant increase (p< 0.01) in firing irregularity (quantified as coefficient of variation (CV) of inter-spike intervals; Fig. 2g) in comparison to controls. On the other hand, DLS-lSN DA neurons recorded with 10 nM monomeric α-synuclein showed no loss of firing frequency or regularity (Fig. 2d, 2f, 2g), in comparison to control solution. In about 65% (n = 9/14) DLS-lSN DA neurons dialyzed with 10 nM α-synuclein fibrils, the spontaneous activity was completely abolished (Fig. 2f) with the membrane potential persistently hyperpolarizing below the threshold for pacemaking (Fig. 2e, 2h). In contrast to neurons exposed to fibrillar α-synuclein, only 25% DLS-lSN neurons (n=3/12) exposed to monomeric α-synuclein ceased their spontaneous activity (Fig. 2f) and showed hyperpolarization of the average membrane potential (Fig. 2h). Indeed, steady state membrane potentials (i.e. membrane average potential in last 10 seconds of recording) were significantly lower in neurons dialyzed with fibrils (p< 0.05) compared to controls and monomers.

As additional controls, we also performed experiments with extracellular exposure to 10 nM α-synuclein fibrils while monitoring firing activity in the on-cell mode. Importantly, we observed that DLS-lSN DA neurons also ceased to fire after local extracellular application of 10nM fibrils but not when control extracellular solution were applied (Fig. S2a, S2b). Also, in on-cell, where the metabolic integrity of the neurons is unperturbed, there was a significant decrease in average firing frequency and increase in CV after extracellular application of fibrils (Fig. S2c, S2d). These experiments essentially rule out that the selective effect of α-synuclein fibrils on DLS-lSN DA was an artifact induced or potentiated by the whole-cell configuration.

Figure 3 shows the firing patterns of mSN DA neurons projecting to dorso-medial striatum in mice. In humans, the most equivalent dorso-medial DA projection would innervate the caudate nucleus, which is less affected in PD. In contrast to the DLS-lSN DA neurons, DMS-mSN DA neurons displayed no loss of pacemaker firing (Fig. 3d, 3f) or membrane hyperpolarization (Fig. 3e, 3h) after dialysis of either 10 nM fibrils or monomers. There was an only about 30% decrease in pacemaker firing frequency after exposure to both fibrillar or monomeric α-synuclein, which was very similar to the about 22% decrease in average firing frequency in control experiments. Some DMS-mSN DA neurons (n=5/11) displayed an increased variability in firing as shown in the trace in figure 3b. This lead to an increase in CV of these neurons, but overall it was not statistically different from the neurons in control group (Fig. 3g). None of the neurons exposed to monomeric α-synuclein displayed any irregularity in pacemaker firing or a decrease in steady state membrane potential. Consistent with their smaller susceptibility during PD, DMS-mSN DA neurons responded – if at all – only weakly to fibrils.

**Figure 3.**
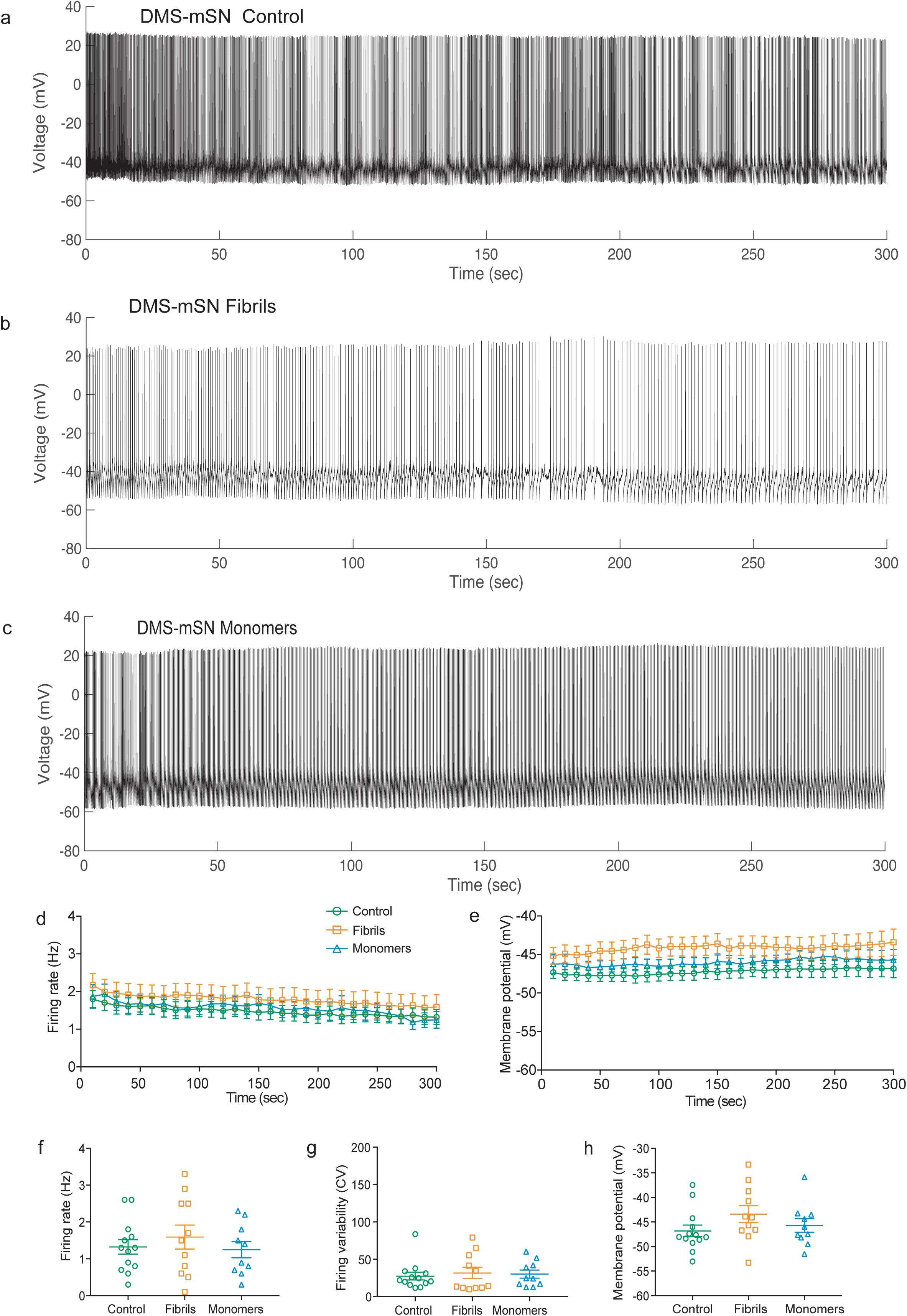
Acute exposure to α-synuclein fibrils does not alter the pacemaker firing of DMS-mSN neurons. Representative trace of whole cell recordings for a duration of 300 seconds from DMS-mSN under **a** control conditions showing regular firing **b** with 10 nM fibrils in the recording solution where mild irregularity **c** with 10 nM monomers in the recording solution showing no change in firing patterns. Graph shows the change in **d** average firing frequency and **e** average membrane potential over the recording duration in various groups. **f** Comparison of steady state firing frequency (i.e. frequency during last 10 seconds of recording) shows no significant change in firing frequency after fibrils or monomers wash-in in comparison to control. **g** Comparison of steady membrane potential (i.e. membrane potential during last 10 seconds of recording) shows no (Ordinary one-way ANOVA followed by Holm Sidak’s multiple comparison test) in membrane potential after fibrils wash-in in comparison to control. All data is represented as mean ± sem. Control- n=13, N=8; Fibrils- n=11, N=4; Monomers- n=10, N=5, where n= number of neurons, N= number of mouse.

Figure 4 shows the firing patterns of VTA DA neurons projecting to ventral striatum, more particularly, the core of nucleus accumbens, which is only mildly affected in PD. There was no significant change in the frequency of firing (Fig. 4d, 4f) or the regularity of firing (Fig. 4g) of these mesolimbic DA neurons in presence of either 10 nM α-synuclein fibrils or monomers. Further, no changes in the average membrane potential were observed in neurons recorded with fibrillar or monomeric α-synuclein (Fig. 4e, 4h). Overall DLS-lSN DA neurons were most severely affected by exposure to fibrillar α-synuclein in comparison to the other two DA subpopulations.

**Figure 4.**
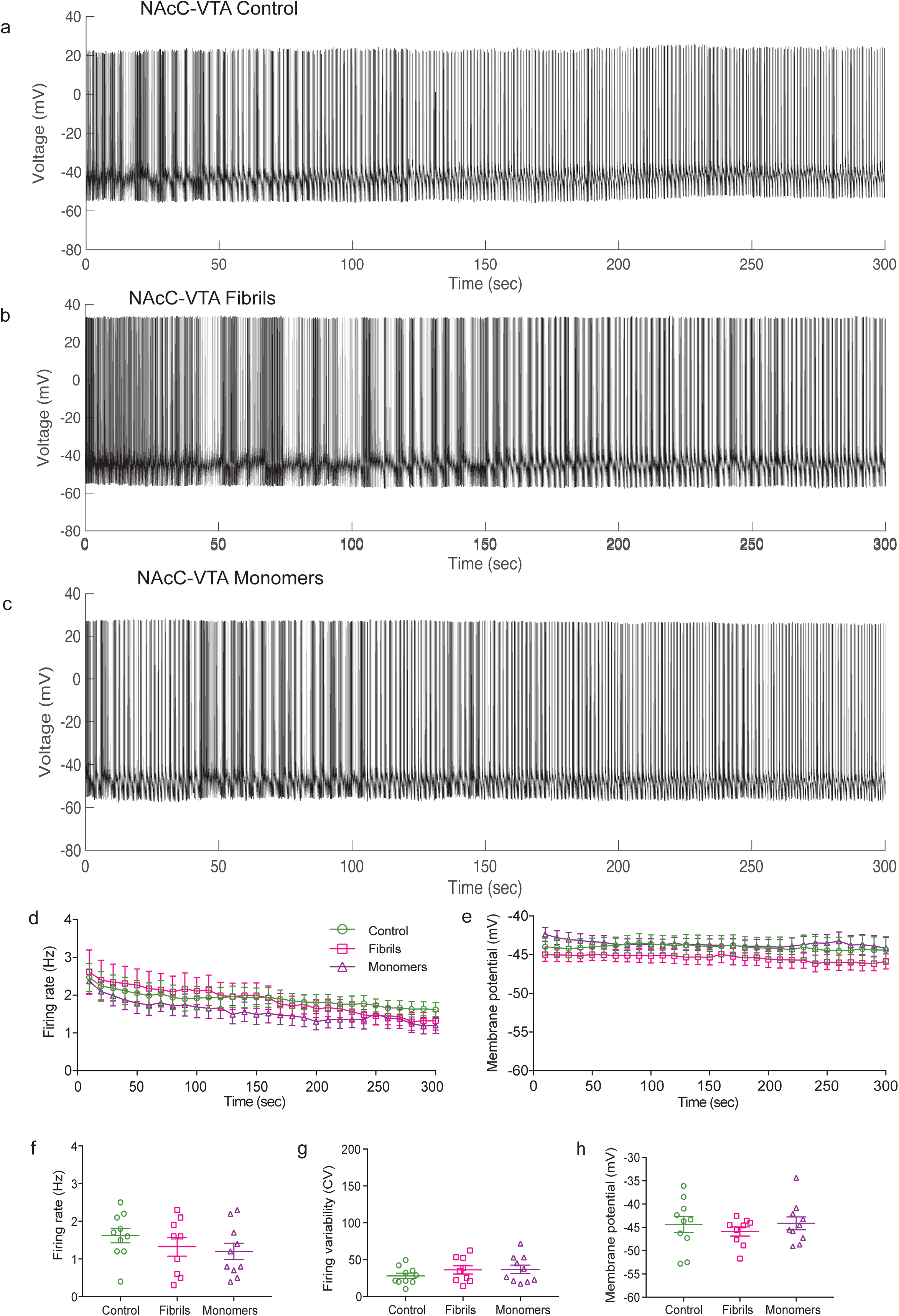
Acute exposure to α-synuclein fibrils does not alter the pacemaker firing of NAcC-VTA neurons. Representative trace of whole cell recordings for a duration of 300 seconds from NAcC-VTA under **a** control conditions showing regular firing **b** with 10 nM fibrils in the recording solution where no irregularity is observed **c** with 10 nM monomers in the recording solution showing no change in firing patterns. Graph shows the change in **d** average firing frequency and **e** average membrane potential over the recording duration in various groups. **e** Comparison of steady state firing frequency and **g** steady state membrane potential show no change (Ordinary one-way ANOVA followed by Holm Sidak’s multiple comparison test) after fibrils or monomers wash-in in comparison to control. All data is represented as mean ± sem. Control- n=10, N=3; Fibrils- n=9, N=3; Monomers- n=10, N=3, where n= number of neurons, N= number of mouse.

### K-ATP channel activation in DA neurons mediates the disruption of firing patterns by fibrillar α-synuclein

Next, we asked whether K-ATP channel activation mediates the α-synuclein fibril induced membrane hyperpolarization and loss of firing activity. We recorded DLS-lSN DA neurons with α-synuclein fibrils in the presence of the K-ATP channel inhibitor glibenclamide (1μM) in the bath solution. Prior to the application of glibenclamide, we observed an 80% decrease in firing frequency as well as membrane hyperpolarization in response to nanomolar α-synuclein fibrils (Fig. 5b, 5d). Similar to previously described changes (Fig. 2), complete cessation of pacemaker firing was observed in around 67% of recorded neurons (n=8/12) leading to a significant decrease in steady state firing frequency in comparison to control (p< 0.001). In contrast, after application of glibenclamide, DLS-lSN DA neurons continued to fire despite the presence of α-synuclein fibrils (Fig. 5c) and showed only 19% decrease in firing rate over the recording period. Steady state firing frequency of these neurons was significantly higher in comparison to the neurons treated with only fibrils (p< 0.001) (Fig. 5f). Some recordings were also obtained with control solution in patch pipette and glibenclamide in the bath (Fig. 5a). In this case, we observed a higher firing frequency (initial frequency around 4.5 Hz) compared to the DLS-lSN neurons recorded regularly with a control solution (initial frequency around 3 Hz) indicating the K-ATP channels were already partially active in this DA subpopulation. This was also supported by on-cell firing frequency data (data not shown). Average firing frequency of glibenclamide-treated control cells (4.8 ± 0.15 Hz, n=11) was significantly higher (p<0.001, unpaired t-test) than the control cells without glibenclamide (3.2 ± 0.3 Hz, n=13).

**Figure 5.**
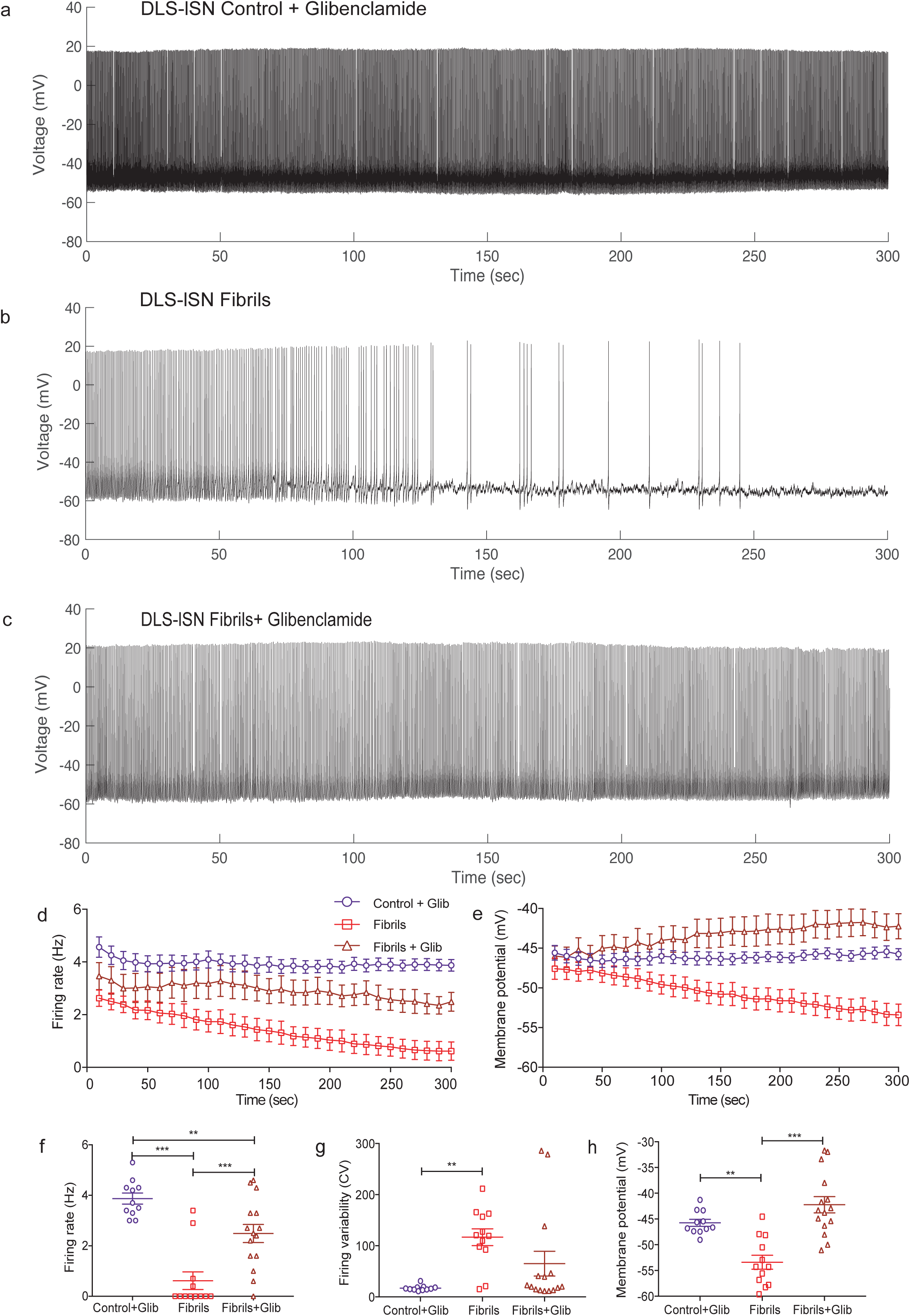
K-ATP channel inhibitor Glibenclamide prevents disruption of firing in DLS-lSN neurons by fibrils. Representative trace of whole cell recordings for a duration of 300 seconds from DLS-lSN neurons **a** with control recording solution and Glibenclamide applied in bath **b** with 10 nM fibrils and **c** with 10 nM fibrils in recording solution in presence of Glibenclamide applied through bath solution. **d** Change in average firing frequency of DLS-lSN neurons in various treatment groups shows prevention of firing loss caused by fibrils after Glib (Glibencamide) treatement. **e** Glib application resulted in an increased average membrane potential even in the presence of fibrils. **f** There was a significant decline in steady state firing frequency (p< 0.001, Ordinary one-way ANOVA followed by Holm Sidak’s multiple comparison test) and **g** increase in firing variability (p< 0.001, Ordinary one-way ANOVA followed by Holm Sidak’s multiple comparison test) in presence of α-synuclein fibrils, which was prevented in presence of Glibenclamide. **h** Wash-in of Glibenclamide also prevents the hyperpolarization of membrane potential observed after α-synuclein fibrils treatment (p< 0.001, Ordinary one-way ANOVA followed by Holm Sidak’s multiple comparison test). All data is represented as mean ± sem. Control+Glib- n=11, N=4; Fibrils- n=12, N=4; Fibrils+Glib- n=15, N=3, where n= number of neurons, N= number of mouse.

Figure 5f compares steady state firing frequencies between the different groups demonstrating significantly higher frequencies in fibrils treated neurons in the presence of glibenclamide as compared to those without glibenclamide (p< 0.001,). Also, fibril-induced increases in CV in comparison to control (p< 0.001) was also not observed after application of glibenclamide. Most importantly, glibenclamide treatment was able to prevent the time dependent membrane hyperpolarization observed in the presence of fibrils (Fig. 5e, 5h). There was a significant increase in steady state average membrane potential in the neurons treated with glibenclamide and fibrils both in comparison to the ones treated with only fibrils (p< 0.01).

In summary, pharmacological experiments identified functional K-ATP channels in DA neurons as a necessary mediator for the α-synuclein fibrils-induced disruptions of pacemaking in DLS-lSN DA neurons.

## DISCUSSION

The exceptionally high susceptibility of DA neurons in the lateral SN to degenerate during PD in comparison to more medial DA SN and VTA neurons has been well-documented (Fearnley and Lees, 1991; Kordower et al., 2013b) but the underlying pathophysiological mechanisms have remained unclear. In essence, it was unknown whether these most vulnerable DA midbrain neurons responded to α-synuclein aggregates in a particular and selective fashion. By studying changes in pacemaker firing patterns of well-defined DLS projecting (*putamen-equivalent in primates)* lSN DA neurons and DMS projecting *(caudate-equivalent in primates)* mSN DA neurons subpopulations, we have now identified differential functional responses of these two DA subpopulations after acute exposure to α-synuclein fibrils. Only the most vulnerable DA neurons in the lateral SN showed acute impairment and even silencing of their cell-autonomous pacemaker activity in response to exposure to α-synuclein fibrils in low nanomolar concentrations. Our work targeted a highly relevant cellular population lost in PD and identified a dramatic pathophysiological response with high selectivity and exquisite sensitivity.

Several previous studies have reported the pathophysiological responses to α-synuclein in DA and non-DA neurons (for review, see (Thakur et al., 2019)). For instance, Wu and colleagues have shown that in the primary hippocampal neurons, the fibrillar form of α-synuclein but not the monomeric form lead to decreased synaptic activity in a time- and dose-dependent manner (Wu et al., 2019). Further, incubation with 500 nM α-synuclein oligomers have also been observed to inhibit long-term potentiation in hippocampal brain slices (Martin et al., 2012; Ferreira et al., 2017). Dialyzing 500 nM α-synuclein oligomers through a patch pipette – a similar approach like in our study - into layer 5 cortical pyramidal neurons induced a drop in input resistance and in turn reduced excitability (Kaufmann et al., 2016). In addition, increase in action potential durations were found to be specific to oligomeric but not monomeric α-synuclein (Kaufmann et al., 2016). However, the relevant ion channels were not identified in these types of studies.

In contrast, we identified here a causal role for K-ATP channels selectively in DLS-lSN DA neurons for fibrillar α-synuclein-induced impairment of pacemaking, a key feature of DA neuronal functioning. We have previously shown that K-ATP channels in DA SN but not DA VTA neurons are also sensitive response elements to nanomolar concentrations of rotenone, a mitochondrial complex I inhibitor (Röper and Ashcroft, 1995; Liss et al., 1999, 2005). Moreover, we showed that the rotenone-induced K-ATP channel activation in DA SN neurons was mediated by mitochondrially generated redox stress (Liss et al. 2005). Although this study did not yet differentiate between medial and lateral DA SN neurons, it might imply that also fibrillar α-synuclein activate K-ATP channels via redox signaling, but more studies are needed to test this possibility explicitly. There is however evidence, that fibrillar α-synuclein increase oxidative stress by several, converging mechanisms including mitochondrial dysfunction and altered calcium handling, potentially also acting on a fast time scale and in the low nanomolar range (Choi et al., 2012; Luth et al., 2014; Deas et al., 2015; Thakur et al., 2019).

Also, our study cannot establish the immediate functional impact of α-synuclein mediated K-ATP channel activation in vivo. While our in vitro data suggest also an acute impairment of DLS-lSN DA neurons (see (Farassat et al., 2019) for functional in vivo properties DLS-lSN DA neurons), our previous in vivo study on K-ATP channels in DA SN neurons suggest that their activation – at least in a certain range - might result in higher excitability and more burst discharge (Schiemann et al., 2012). Again, no selective functional role for K-ATP channels in DA neurons of the lateral SN is known. Finally, we have also demonstrated that global genetic K-ATP channel inactivation dramatically reduced the vulnerability SN DA (but not VTA DA) neurons in a chronic MPTP model (Liss et al. 2005). Assuming a similar outcome for conformer-specific α-synuclein models, K-ATP channel activation might selectively drive neurodegeneration in these most vulnerable DA neurons. As K-ATP channel blockers have been in clinical use for diabetes type II, repurposing for clinical trials in PD would be feasible. However, before this seems warranted, in vivo studies using relevant, chronic fibrillar α-synuclein PD models (Thakur et al., 2017) need to provide evidence that K-ATP channel inhibition – directly by sulfonylureas or more indirectly – e.g. by inhibitors of voltage-gated calcium channels – does indeed reduce selective neurodegeneration and might thus become a stronger candidate for disease-modifying therapies in PD. Finally, if fibrillar α-synuclein does activate K-ATP channels selectively in lateral DA SN neurons also in vivo at some point during disease progression in PD, the – yet unknown – functional consequences could also be potentially useful as an early biomarker of DA system at risk.

## ACKNOWLEDGEMENTS

We thank Jasmine Sonntag and Beatrice Fisher for excellent technical assistance. We are grateful to Josef Shin and Strahinja Stojanovic help with protocols. The study was funded by DFG project grants to JR within CRC1080 and CRC815.

## AUTHOR CONTRIBUTION

PT and JR designed the study and co-wrote the manuscript. PT carried out all experiments and performed data analysis. KL produced and characterized the fibrillar and monomeric α-synuclein.

