## Supplementary material for "Selective K-ATP channel-dependent loss of pacemaking in vulnerable nigrostriatal dopamine neurons by α-synuclein aggregates": Supplemetary information

Figure S1

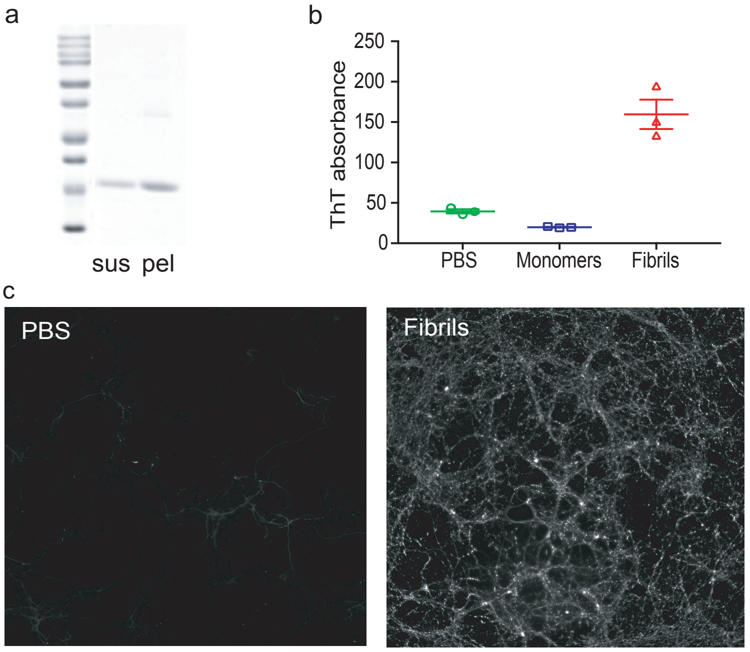


**Figure S1-** Characterization of fibrils. **a** Sedimentation assay-After fibrilization process, reactions were centrifuged (100,000xg, 30 min, 23°C) and both pellet and supernatant were separated on 15% SDS-PAGE gel and visualized with Coomassie Brilliant Blue. Most of the α-synuclein fibrillized and is present in the pellet (pel) fraction while supernatant (sup) contains relatively lower amount of α-synuclein. **b** Fibrillized α-synuclein shows increased absorbance in the Thioflavin T (ThT) assay. Data represents mean ± sem of 3 replicates for each condition. **c** Mouse hippocampal neurons 12 days after transduction with PBS (negative control) or 200 nM fibrils. Immuno-staining for pSer129-α-synuclein revels presence of intraneuronal inclusions.

Figure S2

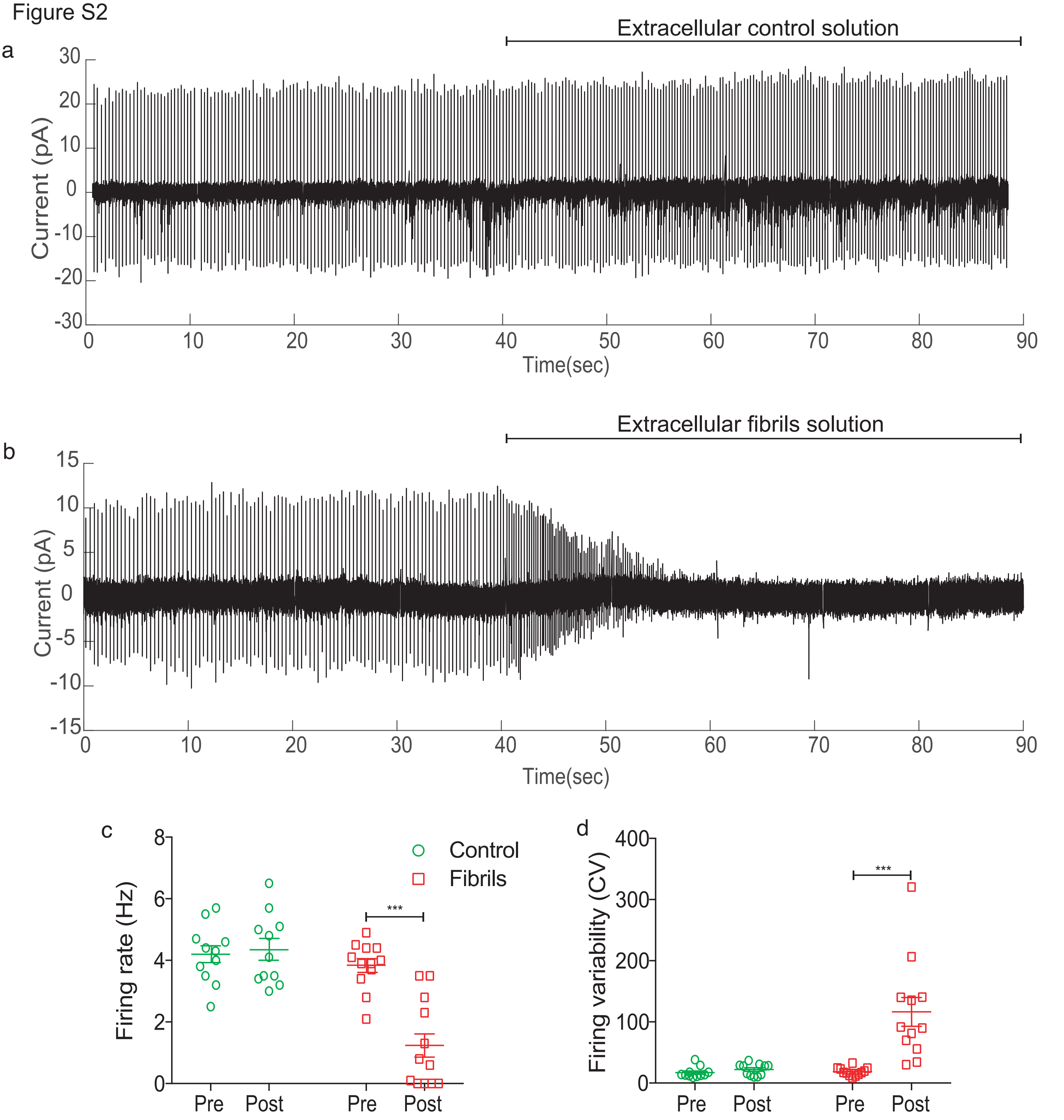


**Figure S2**- Extracellular application of fibrils diminishes the pacemaker firing activity of DLS-ISN neurons. Representative trace of cell-attached (on-cell) recording from DLS-ISN neuron with **a** extracellular control solution puffed on with a second pipette from 40 sec onwards showing no change in firing activity and **b** 10 nM fibrils dissolved in extracellular solution puffed with a second pipette from 40 sec onwards showing a rapid cessation of pacemaker firing. **c** Graph shows the average firing frequency pre and post application of extracellular control solution and fibrils solution. A significant decrease in firing frequency was observed post fibrils application ($p < 0.001$, Ordinary one-way ANOVA followed by Holm Sidak's multiple comparison test) but not post control solution application. **d** Graph shows changes in firing variability (CV) pre and post application of extracellular control solution and fibrils solution. A significant rise in firing variability was observed post fibrils application ($p < 0.001$, Ordinary one-way ANOVA followed by Holm Sidak's multiple comparison test) but not post control solution application. All data is represented as mean ± sem. Control-n=11, N=3; Fibrils- n=12, N=3, where n= number of neurons, N= number of mouse.
